# Somatic drift and rapid loss of heterozygosity suggest small effective population size of stem cells and high somatic mutation rate in asexual planaria

**DOI:** 10.1101/665166

**Authors:** Hosseinali Asgharian, Joseph Dunham, Paul Marjoram, Sergey V. Nuzhdin

**Affiliations:** Department of Biochemistry and Biophysics, School of Medicine, University of California, San Francisco, San Francisco, California, United States of America; Department of Preventive Medicine, Keck School of Medicine, University of Southern California, Los Angeles, California, United States of America; Program in Molecular and Computational Biology, Dornsife College of Letters, Arts, and Sciences, University of Southern California, Los Angeles, California, United States of America

## Abstract

Planarian flatworms have emerged as highly promising models of body regeneration due to the many stem cells scattered through their bodies. Currently, there is no consensus as to the number of stem cells active in each cycle of regeneration or the equality of their relative contributions. We approached this problem with a population genetic model of somatic genetic drift. We modeled the fissiparous life cycle of asexual planarians as an asexual population of cells that goes through repeated events of splitting into two subpopulations followed by population growth to restore the original size. We sampled a pedigree of obligate asexual clones of *Girardia cf. tigrina* at multiple time points encompassing 14 generations. Effective population size of stem cells was inferred from the magnitude of temporal fluctuations in the frequency of somatic variants and under most of the examined scenarios was estimated to be in the range of a few hundreds. Average genomic nucleotide diversity was 0.00398. Assuming neutral evolution and mutation-drift equilibrium, the somatic mutation rate was estimated in the 10^−5^ − 10^−7^ range. Alternatively, we estimated *N*_*e*_ and somatic *μ* from temporal changes in nucleotide diversity *π* without the assumption of equilibrium. This second method suggested even smaller *N*_*e*_ and larger *μ*. A key unknown parameter in our model on which estimates of *N*_*e*_ and *μ* depend is *g*, the ratio of cellular to organismal generations determined by tissue turnover rate. Small effective number of propagating stem cells might contribute to reducing reproductive conflicts in clonal organisms.

## Introduction

Planarian flatworms are a fascinating model system for studying body regeneration. After injury they can reconstruct their body from very small pieces of tissue; and, can grow or “degrow” by regulating the number of cells in their bodies in response to nutrient availability^1,2^. The asexual lines in the clade (including *Girardia tigrina* which we used in our experiments) are generally fissiparous^3,4^: the grown worm splits down the middle creating a head piece and a tail piece; each half is restored to a complete body through positionally regulated cell multiplication, cell death and differentiation^5^. Growth and regeneration rely mainly on a very large number of stem cells (neoblasts) which comprise 25-30% of the cells in a planarian’s body by morphological observation^2,6,7^. Neoblasts are the only dividing cells in Planaria^2^ but BrdU labeling experiments suggest that only a fraction of them are active at any given time^1^. Due to the heterogeneity of morphological and cellular features in stem cells, as well as their partial similarity to early post-mitotic committed progeny cells, mitotic molecular markers such as PCNA and H3P or BrdU labeling cannot visualize stem cells completely and exclusively ^1,8^. Recently, irradiation and transplantation experiments suggested the surface protein Tetraspanin as a reliable marker for isolation of pluripotent stem cells^9^. Nevertheless, methods involving BrdU injection, experimental wounding, or irradiation significantly alter the normal physiology of the animal. Consequently, a quantitative understanding of the number of active stem cells and their relative contributions at each cycle of tissue regeneration during natural growth and reproduction is still lacking.

Planarians are a curious case for evolutionary genetic studies, too. Because asexual planarians do not undergo the single-cell bottleneck of zygote, significant genetic heterogeneity exists within a single worm’s body^10^, which invokes competition among diverged cellular lineages^11^. Mutator alleles remain linked with the mutations they cause in these non-recombining genomes and somatic mutations are transmitted to future generations. Theoretically, deleterious mutations (if not completely recessive) could be eliminated at the cellular level locally before they reach a frequency that can affect organismal fitness, while beneficial mutations give the mutator lineage a competitive advantage. This dynamic predicts a higher optimal somatic mutation rate in clonal organisms^12^.

In this study we present a simplified population genetic model of the life cycle of an asexual planarian. Most of what is known about planarian biology comes from the study of *Schmidtea mediterranea* ^13–15^. In this study we focused on a less well understood species *Girardia cf. tigrina.* In our model, somatic cells play the role of asexually reproducing individuals in an expanding population, which is the planarian body. Over time, this population doubles in size and splits into two subpopulations, i.e., the head and tail pieces. We showed that the allele frequency spectrum of somatic variants in our model system is shaped more strongly by genetic drift (random fluctuations affected by population size) versus genetic draft (selection acting on tightly linked loci). Then, we proceeded to estimate somatic mutation rate from *N*_*e,sc*_ and the observed nucleotide diversity (*π*) according to expectations of the neutral theory. We tracked fluctuations in the frequency of somatic variants and average nucleotide diversity over >10 generations and applied the theories of genetic drift and mutation-drift interaction at the cellular level to estimate the effective size of the stem cell population *N*_*e,sc*_ and the somatic mutation rate *μ*_*som*_.

## Results

A line of lab-reared asexual flatworms was established from a single individual. Cytochrome oxidase subunit 1 (COI) DNA barcode and proportion of total reads mapping to different Dugesiidae genomes both identified this lineage as *Girardia cf. tigrina* (Tables S1 and S2). In several years of maintaining this line, no instance of sexual reproduction was observed. For this study, the worm lineage was followed for 14 generations of splitting and regrowth. Genomic libraries were prepared from tail pieces after splits 2 (II), 6 (VI), 8 (VIII), 10 (X), 12 (XII) and 14 (XIV) and sequenced in 2 or 3 replicates (Fig. 1). One of the replicates from sample XIV failed, making samples II and XII the two samples farthest apart in time that had replicated sequenced libraries. The fastq files were trimmed, filtered and deduplicated using Trimmomatic and BBTool’s Clumpify and evaluated by Fastqc (Table S3). FreeBayes was run on individual replicates as separate samples first. Bi-allelic SNPs with coverage 10-40X in all samples were subjected to principal component analysis (PCA). PCA confirmed that the difference between replicates was indeed much smaller than that between biological samples (Fig. S1). FreeBayes was run a second time merging all the replicates of each biological sample. Allele frequencies of positions with coverage 10-60X from the merged-rep VCF were used for population genetic analysis. Mean sequencing coverage of SNPs was 12.1X for sample XIV and 19.9-36.6X for the other merged-rep samples.

**Figure 1.**
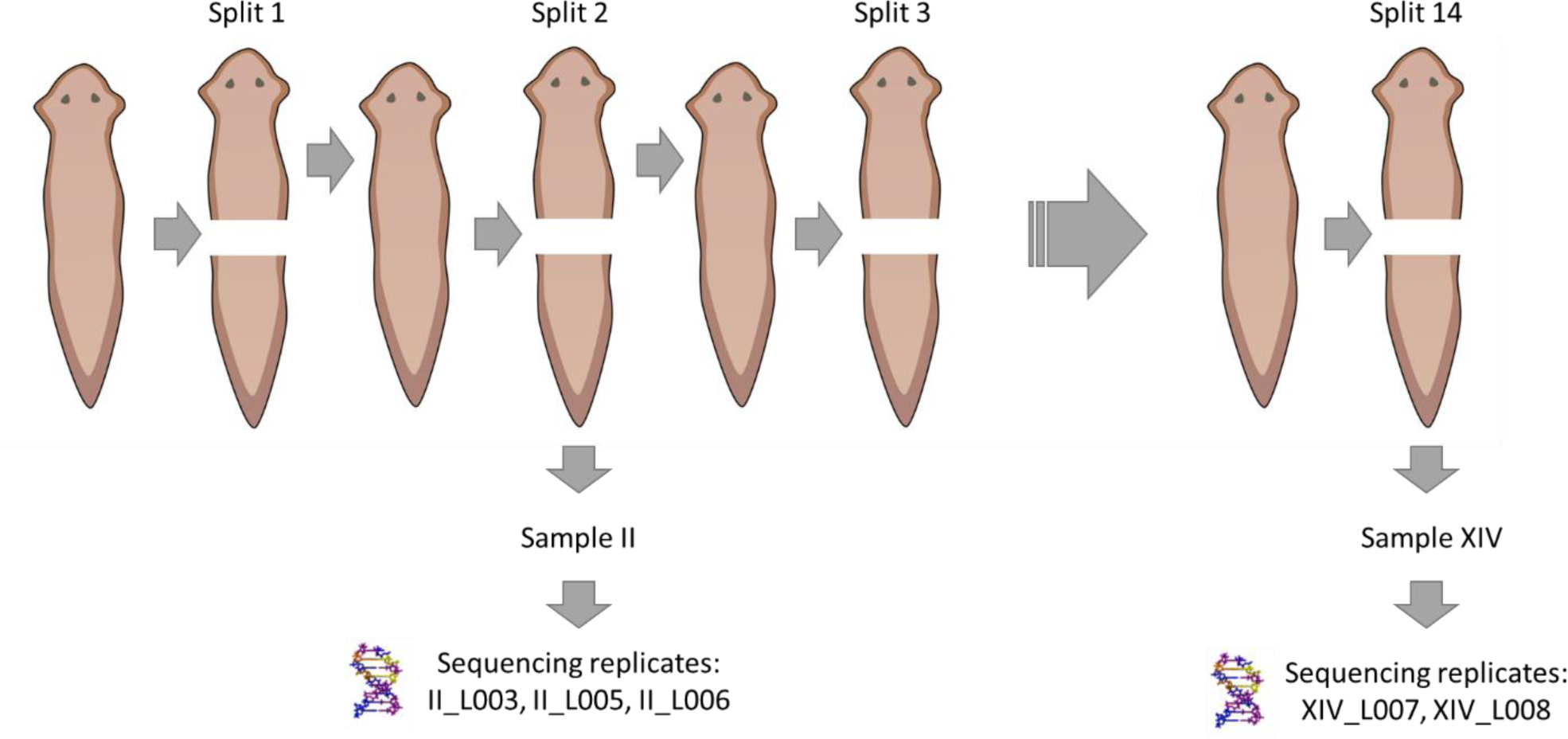
Experimental design of the project. Sample names have two parts: the first part is the split after which the biological sample was derived i.e. organismal generation; the second part is the lane on the HiSeq machine where that replicate was sequenced. For example, sample II_L003 or II.3 was derived after split 2 (generation 2) and sequenced on lane 3. Sample XIV.8 was disqualified due to the loss of its forward sequencing fastq file effectively making sample XII (generation 12) the latest generation that was sequenced in replicates (XII.5 and XII.6).

**Table 1.** Estimates of stem cell effective population size *N*_*e,sc*_ and somatic mutation rate *μ*_*som*_ inferred from temporal variation of allele frequencies and loss of heterozygosity. Samples from the II-XII and II-XIV generation intervals have been analyzed according to scenarios of cell immortality, or various rates of tissue turnover (*g* = 0.5, 1, 5, 10, respectively). *N*_*e*_ calculations were based on 370 SNPs selected for having AAF 0.1-0.9 at the start of the experiment (generation II) and being covered 10-60X in all merged-rep samples. When estimating *N*_*e*_ and *μ* from loss of heterozygosity rate, *π* from all generations (with possible exclusion of generation XIV) were used, not just the first and last generation. *g*: ratio of cellular / organismal generations, *F̄*_*k*_: arithmetic average of *F*_*k*_over loci, *S*_*i*_ : harmonic mean of sequencing coverage of loci at the initial sampling point (generation II), *S*_*j*_: harmonic mean of sequencing coverage of loci at the final sampling point (generation XII or XIV as indicated in the first column), *μ*:_*som*_ : expected somatic mutation rate under mutation-drift equilibrium, *π*_0_: heterozygosity extrapolated to generation zero (two generations before the start of sequencing). - : the nonlinear least square algorithm did not converge after 100 iterations.

| Gen.s | $g$ | $\Delta t_{cell}$ | Estimated from somatic drift | | | | Estimated from loss of heterozygosity | | |
| --- | --- | --- | --- | --- | --- | --- | --- | --- | --- |
| | | | $\bar{F}_k$ | $S_i, S_j$ | $N_{e,sc}$ | $\tilde{\mu}_{som}$ | $\pi_0$ | $N_{e,sc}$ | $\mu_{som}$ |
| II-XII | 0.5 | 5 | 0.1433 | 31.25, 21.73 | 71.10 | 2.7989E-05 | - | - | - |
| II-XIV | 0.5 | 6 | 0.1378 | 31.25, 15.14 | 138.49 | 1.4369E-05 | 0.0048 | 5.19 | 3.14E-04 |
| II-XII | 1 | 10 | 0.1433 | 31.25, 21.73 | 141.70 | 1.4044E-05 | - | - | - |
| II-XIV | 1 | 12 | 0.1378 | 31.25, 15.14 | 276.48 | 7.1976E-06 | - | - | - |
| II-XII | 5 | 50 | 0.1433 | 31.25, 21.73 | 706.52 | 2.8166E-06 | - | - | - |
| II-XIV | 5 | 60 | 0.1378 | 31.25, 15.14 | 1380.39 | 1.4416E-06 | 0.0048 | 51.9 | 3.14E-05 |
| II-XII | 10 | 100 | 0.1433 | 31.25, 21.73 | 1412.54 | 1.4088E-06 | 0.0047 | 478 | 0 |
| II-XIV | 10 | 120 | 0.1378 | 31.25, 15.14 | 2760.27 | 7.2094E-07 | 0.0048 | 104 | 1.57E-05 |

### Divergence from the reference genome

In the merged samples, divergence was calculated as the ratio of the number of positions in the coverage 10-60X range with *AAF* > 0.99 to the total number of positions covered 10-60X. Average divergence from the six samples was 1.112% (range 1.03-1.28%) which corroborates taxonomic identification of our specimens as conspecific or congeneric to *Girardia tigrina*.

### Somatic drift vs. somatic draft

The allele frequency spectra from samples II and XII are plotted in Fig. S2. Assuming the reference allele to be ancestral in most positions, patterns of the alternative (derived) allele frequency roughly match the neutral expectation except for a local peak of *f*_*alt*_ ≅ 1 representing positions of fixed divergence from the reference. Simulations have shown that in the absence of recombination, strong selection on linked loci can distort the distribution of allele frequencies in a specific way: the density of derived allele frequency falls off much more steeply under linked selection (draft) than under neutral evolution (drift)^16^. Denoting derived allele frequency *ν* and the corresponding density *f*(*ν*), derived allele frequency falls as *f*(*ν*)∼*ν*^−1^ under drift but as *f*(*ν*)∼*ν*^−2^ under draft^16^. In our dataset, the *f*(*ν*)∼*ν*^−1^ model (*R*^2^ = 0.938) fits the data better than the *f*(*ν*)∼*ν*^−2^ model (*R*^2^ = 0.832) (Table S4 and Fig. S3). This suggests that: 1) The observed AFS is more consistent with drift than draft, although draft cannot be rejected and probably plays some role (0.832 is not much smaller than 0.938). 2) The drift model fits quite well, which validates the idea of estimating stem cell *N*_*e*_ based on somatic drift. Fig. S4 illustrates temporal fluctuation in alternative allele frequency of 9 randomly picked loci.

### Effective size of stem cell population and somatic mutation rate

We calculated *N*_*e,sc*_ and somatic mutation rate *μ*_*som*_ by two different approaches: 1) using somatic drift and the magnitude of change in somatic variant frequencies (equations 1-3, see Methods), and 2) using the rate of change in heterozygosity (*π*) across generations (equation 7). In both cases, the parameters of interest were estimated with and without generation XIV which was not sequenced in replicates (Table 1). The first approach uses allele frequencies from samples at the two ends of the longest interval of generations, II-XII or II-XIV. The second approach uses nucleotide diversity (*π*) from all generations. We estimated *F̄*_*k*_ and *N*_*e*_ from individual replicates of generations II and XII to evaluate replicate reproducibility (Table S5). Subsampling of SNPs showed that although the 370 SNPs we analyzed were all located on the same contig and were inherited together during asexual reproduction, they provided relatively independent pieces of information on somatic drift (Fig. S5). This was probably because we sequenced tens of thousands of cells but were looking at positions with coverage 10-60X. Reads mapping to different positions likely came from different, possibly divergent cell lineages having had different past trajectories and accumulated different mutations and gene conversions through generations of clonal reproduction. This was equivalent of sequencing multiple loci in different subsets of individuals in the population. The estimates of *N*_*e,sc*_ in Table 1 change considerably depending on the choice of final sampling generation (XII or XIV). This more than expected due to somatic drift alone. It is possible that there is more biological stochasticity in stem cell division patterns over time depending on environmental factors that were not accounted for in our experiments. It could also be influenced by experimental errors which suggests that future experiments must involve more control conditions, more replicated and more time points spaced over longer intervals. *N*_*e,sc*_ estimate from drift analysis also varies almost linearly with the assumed number of elapsed generations (*t*), which in turn depends on the ratio of cell to organismal generations (*g*). Although the rate of tissue turnover in planarians under natural physiological conditions has not been measured experimentally, a theoretical lower bound for *g* can be obtained by assuming that 1) a stem cell’s proliferation rate is independent of its age, and 2) when the worm grows fast under favorable conditions, most cells are young, and therefore the rate of apoptosis is negligible compared to the rate of cell division. Under these assumptions, the number of cellular generations will be approximately half the number of organismal generations; because, after each split and regrowth event, half the cells come directly from the previous generation (number of generations elapsed = 0) and the other half are newly produced by stem cells (number of generations elapsed = 1), bringing the average over all cells to 0.5. Although this assumption is not realistic, it offers a way to estimate min (*N*_*e,sc*_) and max (*μ*_*som*_). Homeostatic tissue turnover becomes increasingly significant the more slowly the worm grows because more and more somatic cells age, die, and are replaced, adding to the number of elapsed cellular generations. The smallest estimate from drift analysis is *N*_*e,sc*_ = 71.1 which corresponds to *g* = 0.5 based on allele frequencies from generations II and XII (Table 1).

Nucleotide diversity (*π*) for each sample was estimated as the product of nucleotide diversity at bi-allelic SNPs (*π*^∗^) and the proportion of polymorphic positions in the sample. Average nucleotide diversity across all samples was *π̅* = 0.00398. Assuming mutation-drift balance under neutrality, the equilibrium somatic mutation rate was calculated according to the haploid form of the equation *π* = 2*N*_*e*_*μ* or 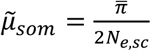. Estimates of *N*_*e,sc*_ and *μ*: *som* under several scenarios are given in Table 1. The eight examined parameter sets estimate *μ* in the 2.7 × 10^−5^ − 7.2 × 10^−7^ range. These values should be interpreted with caution because nucleotide diversity shows a decreasing trend down the generations in our experiment (Fig. 2) indicating that the system may not be at drift-mutation balance and somatic genetic variation may not be in a steady state.

**Figure 2.**
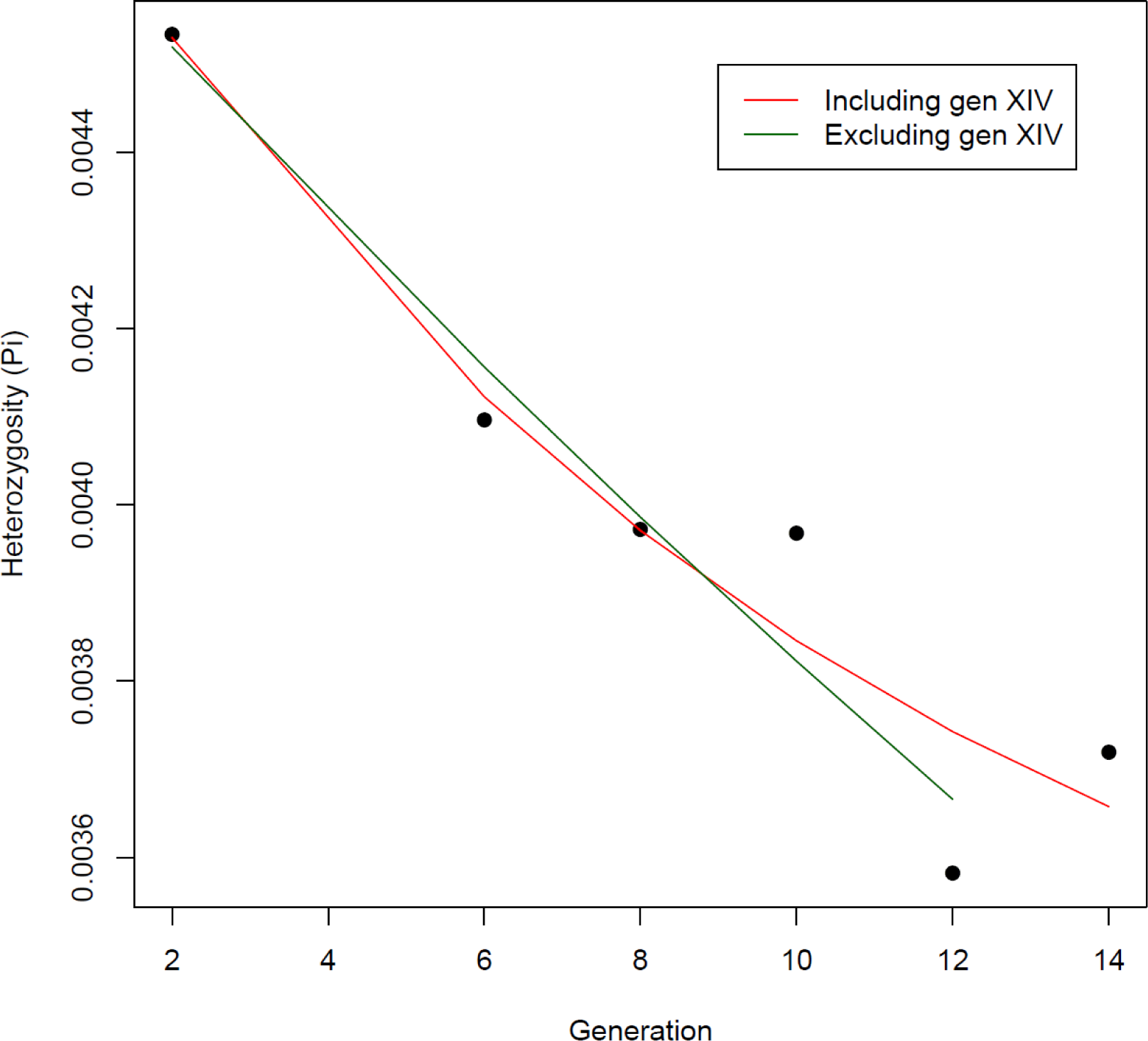
Decreasing trend of nucleotide diversity over generations. The best nonlinear least square fits −(^1^ +2*μ*)*t* to equation 7, 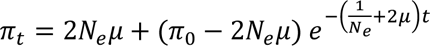, are plotted over the observed values (black circles). Different *g* values result in correspondingly scaled estimates of *N*_*e*_ and *μ* (Table 1) but the same predicted values of *π*.

We derived a model describing the change in heterozygosity (*H*) with combined effects of drift and mutation in its general form i.e. without the assumption of equilibrium (Eq. 7). According to this equation, the difference between the current value of *H* and its equilibrium value is reduced by an exponential factor of 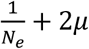 per generation. Equation 7 cannot be linearized and there is no closed form solution to find the best fit; thus, numerical methods must be used. We used a nonlinear least square algorithm to find parameter values for *N*_*e,sc*_, *μ*_*som*_ and *π*_0_ that best fit the observed trend of *π* over generations (Fig. 2, Table 1). These are different from estimates inferred from somatic drift and temporal variance of allele frequencies and suggest even smaller *N*_*e*_ and larger *μ*_*som*_.

## Discussion

In the absence of direct molecular and cellular evidence and effective tools for *in vivo* analysis of natural growth and regeneration in planarians, we decided to use the classical population genetic theory to model stem-cell-based body regeneration in an asexual line of *G. tigrina*. Natural populations of *G. tigrina* comprise sexual and asexual lines and show variable ploidy (2n or 3n)^3,17^. Visual inspection of somatic allele frequencies of nine example loci showed what appeared to be random fluctuations (Fig. S4). Evaluating the density of the allele frequency spectrum further corroborated the role of somatic drift at the cellular level (Table S4, Fig. S3).

The properties of our experimental systems match the underlying assumptions and structure of the theoretical model reasonably well. In the original scheme theorized by Waples, a population is sampled at two time points and an estimate of effective population size (*N*_*e*_) is derived from observed temporal changes in allele frequencies^18^. To make the model more analytically tractable and free of certain restricting assumptions, they recommend a sampling before reproduction plan to ensure that there will be no overlap (and therefore no covariance) between the reproducing individuals (contributing to *N*_*e*_) and individuals sampled for allele frequency estimation. Such a separation is guaranteed in our model system as tail pieces are sequenced while head pieces grow to create the next generation. The theory we used assumes discrete generations, but we know planarian cells do not divide and die synchronously. Results from applying the classical *N*_*e*,*Fk*_ method to populations with overlapping generations may be biased^19^. Accuracy of *N*_*e*_ estimation can be improved by examining more loci, using loci with intermediate allele frequencies, sampling more individuals and increasing the time between samples (at least 3-5 generations apart, preferably more)^20,21^. Following these guidelines, we estimated *N*_*e,sc*_ from the longest interval between our samples. Our estimate of *N*_*e,sc*_ comes from 370 positions which is much larger than recommended number of loci to achieve acceptable accuracy and precision in the references. We also omitted SNPs with minor allele frequency <0.1. Subsampling analysis confirmed that despite being on the same contig, these positions provided independent bits of information (Fig. S5).

Our estimates of the effective number of stem cells even under the most permissive scenarios are still much smaller than the number of stem cells suggested by microscopic observations (tens of thousands). It has been suggested that 20-30% of cells in a worm’s body are stem cells^22–24^. A 1-cm long Dugesia was estimated to contain approximately 2 × 10^5^ cells^25^. This gives an estimate of 40000-60000 stem cells which is 1 or 2 orders of magnitude higher than our estimate of a few hundreds to a few thousands (Table 1). The most likely explanation is that a small fraction of stem cells, e.g., stem cells closer to the fission site, contribute disproportionately highly to regeneration. It should be noted that we estimated the number of active stem cells in the head piece, which is known to have fewer stem cells than the tail piece, with the area anterior to the eyes practically devoid of any stem cells. Underestimation of homeostatic tissue turnover rate reflected in *g* (ratio of cellular to organismal generations) is a possibility. Hopefully, experimental data in the future will quantify tissue turnover and *g* can be specified with more certainty. It has been shown that selection will not affect *F*^^^_*k*_ much under plausible *t* circumstances, especially when 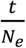 is small^26^. It has been suggested that variable selection and changes in demographical parameters can lead to overestimation of *F*_*k*_ and underestimation of *N*_*e*_^18^; but these are unlikely to have played significant roles in our model system of worms reared in controlled lab conditions. Effective population of stem cells can vary from generation to generation not only due to stochasticity of activation at the cellular level, but also because worms do not always split exactly equal or consistently proportionate pieces. The number of stem cells in the head piece will depend on the size of the head piece. When the population size (of stem cells or otherwise) varies from generation to generation, *N*_*e*_ estimates the harmonic mean of *N*s across generations which is influenced very strongly by the smallest values of *N*.

One of the caveats of our models was that we used a well-mixed model equivalent of a panmictic population. Existence of body structure in planaria means that this assumption is not accurate. However, this is a common simplifying assumption in almost all population genetic models except those focusing specifically on spatial heterogeneity. It is also assumed in the theory we used here for its original purpose of estimating *N*_*e*_ in organismal populations, although natural populations of animals and plants are not realistically panmictic. The planarian body structure may cause *N*_*e,sc*_ to be smaller than the microscopically observed number of stem cells, since stem cells closer to the fission site probably play a more important role in regeneration. However, it is unlikely to violate the assumptions of our model and affect its conclusions substantially because of three mitigating factors: 1) After experimental amputation, natural fission, or during the processes of growth and degrowth, the worm’s body undergoes extensive reshaping (known as morphallaxis) ^2,28^. This means that body structure is likely not preserved from generation to generation, and therefore random activation of stem cells is an appropriate model for estimation of effective population size. 2) Irradiation-amputation experiments show that stem cells or their progeny can migrate long distances from their original position to the wound site to contribute to regeneration ^1,2^. The reconstruction of the body after fission is not restricted to stem cells adjacent to the fission site although they might contribute more. 3) It is noteworthy that population structure (lack of complete mixing) has a similar effect on probability of identity by descent of alleles (inbreeding coefficient, denoted *F*) which is the basis for the definition of *N*_*e*_: the effective population size of a real population is equal to the consensus size of an ideal population which results in the same amount of identity by descent or *F* per generation. Strong population structure reduces heterozygosity because it effectively divides the actual population to a number of small drift-prone sub-populations with poorly connected gene pools. In the case of our experimental system, population structure within the stem cell population would mean that not all stem cells are equally likely to contribute to regeneration, with the ones close to the fissure being more likely to do so. Therefore, the main conclusion of small *N*_*e*_ remains valid, but it could be interpreted as population structure or random activation of a small fraction of stem cells. However, population geneticists do not solely attribute *N*_*e*_ to neutral processes; there is abundant literature on the effects of background selection and selection sweeps on *N*_*e*_ ^29,30^. It was therefore important to determine which is the primary force shaping *N*_*e*_ in our system: drift or selection? Drift inevitably happens in any population of finite size, whereas selection only sometimes occurs with an effective strength. Drift is the default assumption unless there is significant evidence that it cannot explain the data in which case other processes are evoked. In our analysis of allele frequency spectrum, we found no evidence to believe that selection is a stronger force than drift in this dataset. We are not dismissing the possible occurrence of selection, but the data show that it is not a significant determinant of *N*_*e*_ in our system. If selection is not strong enough to alter the AFS significantly, to the point that AFS is slightly better explained by drift, using a more complex model incorporating weak selection does not seem warranted.

Another limitation of this study is that the value of *g*, the ratio of cellular to organismal generations, is unknown. As far as we know, it is yet unknown even for the most well-studied planarians. Our estimates of *N*_*e*_ and *μ* from both methods depend on this parameter. Our model is still useful because it provides a relatively simple analytical framework for the estimation of two important cellular and genetic parameters from sequencing data, and reduces it to one unknown value. The value of *g* depends on several aspects of planarian biology, most importantly the rate of homeostatic cell turnover rate. Future experiments will focus on quantifying this value. In the absence of *g*’s precise value, we decided to determine its upper and or lower boundaries and calculate *N*_*e*_ and *μ* based on those boundaries. The upper bound for *g* is theoretically infinite and practically depends on environmental factors e.g. the feeding regime as well as inherent ones. The lower bound for *g*, however, is 0.5, which would be the case if all cells were immortal. Although we can be certain that this is not a realistic possibility, it provides a useful basis for calculation of min (*N*_*e*_) and max (*μ*).

The estimated somatic mutation rate is of the order of 10^−5^ − 10^−7^ which is orders of magnitude higher than the norm in sexually reproducing eukaryotes (often in the 10^−8^ − 10^−9^ range). There are several points of consideration here. First, in most organisms, including humans, the rate of mutation in somatic tissues is about an order of magnitude higher than in the germline^31,32^. Second, theory predicts that mutation rate evolves to a minimum rate in sexual populations but could evolve to a non-minimal optimum in asexual populations under particular circumstances^33,34^. It has been shown that in the presence of strong selection among somatic cell lineages, a higher optimal mutation rate can evolve, because mutator alleles can benefit from the advantageous mutations they cause while deleterious mutations are eliminated before they get the chance to be transmitted to next generation^11,12^. The high level of somatic heterogeneity, also observed by Nishimura et al.^10^, can provide the basis for such strong somatic selection. Third, the neutral expectation of *π* = 2*N*_*e*_*μ* is derived assuming a steady state level of genetic variation (mutation-drift balance). However, we observed a rapid loss of heterozygosity in our samples over the generations (Fig. 2), which is in contrast with previous observations in a sexual line of *S. mediterranea* ^27^. Inference from loss of heterozygosity produces even smaller *N*_*e*_ and larger *μ*_*som*_ estimates than those estimated from drift. Most natural systems are expected to maintain a steady state level of genetic variation over long term. We speculate that the observed decreasing trend in *π* is partly due to the fast growth and splitting of worms under favorable lab conditions. Under slower growth, more homeostatic tissue turnover would happen which would allow for the accumulation of as many new mutations as eliminated by somatic drift. Spontaneous sexualization has been reported in some G. tigrina lines which could potentially restore genetic diversity^7^ although this was not observed during our experiments.

Although the estimates from both methods change in the same direction with the key parameter *g*, there is considerable difference between the two sets of estimates. This will be addressed in future work by gathering more data and developing more complex simulation-based models incorporating body structure, homeostatic tissue turnover, variable growth rates and splitting patterns.

## Methods

### Worm collection and maintenance

The worms were collected from a stream in Almese, Italy on September 23, 2009. They were separated into individual vials and kept in standard rearing conditions and fed beef liver once a week followed by water exchanges. Over more than two years, the worms reproduced exclusively asexually through fissipary. All the splits occurred naturally; no artificial cutting, wounding or injection was performed. At the beginning of the experiment, a single lineage was followed for 14 generations of natural splitting and regrowth. After generations II, VI, VIII, X, XII and XIV, the tail piece was frozen for sequencing while the head piece was left to grow and further the lineage (Fig. 1).

### Sequencing, QC and duplicate removal

Genomic libraries were produced from frozen tail pieces with 2-3 replicates according to the protocol described previously^35^ and sequenced on Illumina HiSeq. Raw fastq files were examined by FastQC (https://www.bioinformatics.babraham.ac.uk/projects/fastqc). Low quality segments were removed and short trimmed reads (<36 bases) dropped using Trimmomatic v.0.38^36^ in the paired-end mode with the following options: */<PATH>/trimmomatic-0.38.jar PE <INPUT files> <OUTPUT files> ILLUMINACLIP:/<PATH>/adapters/TruSeq3-SE.fa:2:30:12:1:true LEADING:3 TRAILING:3 MAXINFO:40:0.4 MINLEN:36*. Only proper pairs (both mates surviving) were processed further. PCR duplicates were removed using the Clumpify command of BBTools (https://jgi.doe.gov/data-and-tools/bbtools) in paired-end mode with the following options: */<PATH>/clumpify.sh -Xmx230g <INPUT files> <OUTPUT files> dedupe=t reorder*. In addition to removing duplicates, Clumpify sorts fastq files for more efficient compression which reduces compressed file size and accelerates future processing steps. Trimmomatic and Clumpify output files were re-evaluated by FastQC.

### Taxonomic identification

Based on initial morphological inspection, the specimens were tentatively assigned to family Dugesiidae. Molecular identification was carried out using cytochrome oxidase subunit 1 (COI) barcode sequences ^37^. Pre-aligned COI barcodes of 72 species belonging to the order Tricladida including 16 species from family Dugesiidae were downloaded from the Barcode of Life database (boldsystems.org) public records. Whole genomic sequencing reads from two of our samples were mapped (separately) to the Tricladida COI sequences using bowtie2 (--end-to-end alignment default options). All our samples come from a single founder and belong to the same lineage; however, two samples were tested to ensure the reproducibility of identification. To verify the COI identification by a wider genomic scan, 1000 reads were randomly subsampled from each one of the 24 fastq files pertaining to the 12 paired-end sequenced samples and then pooled together. This pooled fastq was mapped to the concatenated fasta file comprising the four published Dugesiidae genomes on NCBI (*Schmidtea mediterranea* asexual strain CIW4, *S. mediterranea* sexual strain S2, *Dugesia japonica* and *Girardia tigrina*). Reads were mapped using bowtie2 end-to-end and local alignment options (both in the --very-sensitive mode). Although the samples had been sequenced paired-end, for taxonomic identification purposes alignments were performed separately for forward and reverse reads in the unpaired mode to avoid under-mapping of pairs due to insert size variation caused by indels and structural variations, which would not be uncommon in the taxonomic range of a large and diverse order such as Tricladida.

### Mapping and Variant calling for population genetic analysis

Proper trimmed and deduplicated pairs were aligned to the *G. tigrina* genome using *bwa mem* with default options except setting the minimum seed length to 15 (*-k 15*) to facilitate the mapping of more divergent reads^38^. Variants were called using FreeBayes^39^ which is specifically suited to variant calling from Pool-seq data^38^ and has been used to study comparative genomics of flatworms before^40^. FreeBayes was run with options *-F 0.01 -C 1 -m 20 -q 20 --pooled-continuous --use-reference-allele*. This configuration is recommended for variant calling from pooled-seq data with an unknown number of pooled samples (https://github.com/ekg/freebayes) (in our study, the exact number of cells per sample in unknown). The *--use-reference-allele* option ensures that the output includes positions where the base call in the samples differs from the reference allele even if they are monomorphic across the samples. Reads that might map to several parts of the genome were discarded by applying the -m 20 filter (minimum mapping quality of 20). This is especially important in working with potentially highly repetitive genomes which is the case for well-studied flatworms^41^. The current *G. tigrina* assembly comprises >255k scaffolds. This created an error in the running of FreeBayes which we suspect was due to an internal algorithmic step concatenating the names of all chromosomes into a single string. To circumvent this problem, and since we needed only tens of SNPs for a reliable estimate of *N*_*e*_, the bam files and the reference fasta files were filtered to contain only the longest contig (MQRK01218062.1, length=267531). The bam files were manually re-headered in two ways: 1. Rep-by-rep: A different sample name (bam header ‘SM’) was assigned to each replicate 2. Merged-reps: The same sample name (bam header ‘SM’) was assigned to all replicates of the same biological sample, the effect of which is to pool the reads from replicates during variant calling and the consequent calculation of allele frequencies. Sample XIV.8, for which only the reverse mate sequences were available, was excluded from variant calling and later analyses. Thus, all remaining biological samples except sample XIV were represented by at least two independently sequenced replicates. Coverage statistics of the re-headered bam files were obtained using the *samtools mpileup* function. VCF files were filtered using *vcftools* to keep bi-allelic SNPs only. A custom python code was run to keep only SNPs in the coverage range of 10-40X and 10-60X in rep-by-rep and merged-rep VCFs, respectively. Only samples from generations II and XII were used for population genetic analyses (see Results). Consistency of allele frequency estimates among replicates was evaluated via principal component analysis.

### Population genetic analyses

Divergence from the reference genome was calculated from each merged-rep sample as the ratio of SNPs covered 10-60X with alternative allele frequency *AAF* > 0.99 to the total number of positions covered 10-60X in the corresponding sample.

To compare the influence of drift and draft on the allele frequency spectrum, a vector of densities of allele frequencies for SNPs with *AAF* 0.01 − 0.9 was obtained using the density() function in R. Denoting *AAF* as *ν* and its corresponding density as *f*(*ν*), the goodness-of-fit of three linear models *f*(*ν*)∼*ν*, *f*(*ν*)∼*ν*^−1^ and *f*(*ν*)∼*ν*^−2^ were tested by their p-values and *R*^2 16^.

Effective population size of stem cells was calculated based on the change in allele frequencies of SNPs between generations using the theory laid out by Waples^18,42^. In this method, a parameter *F*_*k*_ is calculated from the transgenerational difference in allele frequencies the expected value of which depends on sample sizes at the initial and final sampling points (*S*_*i*_ and *S*_*j*_), number of generations elapsed (*t*) and effective population size (*N*_*e*_). We calculated *F*_*k*_ at each SNP as:

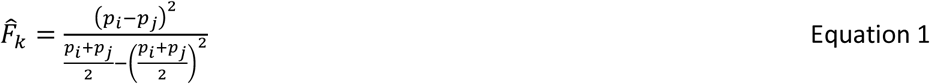

Where *p*_*i*_ and *p*_*j*_ are the frequency of the alternative allele at generations *i* and *j*, respectively (here: *i* = 2, *j* = 12). Only SNPs with *AAF* 0.1 − 0.9 were included to avoid biases introduced by very rare minor alleles^18^. The average *F*_*k*_over loci (*F̄*_*k*_) was plugged into the following equation to obtain *N*_*e*,*sc*_ :

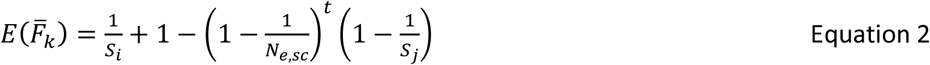

Equation 2 can be rearranged to obtain *N*_*e,sc*_ :

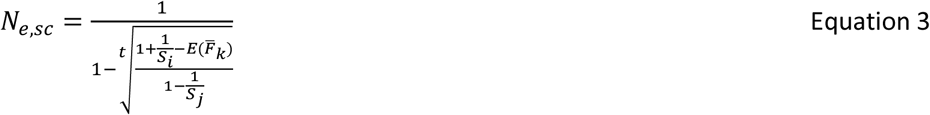

In the original formulation designed for a single locus, *S*_*i*_ and *S*_*j*_ are sample sizes at generations *i* and *j*^18^. Here, sample size is replaced by sequencing coverage. *S*_*i*_ and *S*_*j*_ are harmonic means of sequencing coverage across all examined SNPs in generations *i* and *j*, respectively. The number of generations elapsed between the two sampling points is represented by *t*. Since we are modeling cells as individuals, *t* in our calculations must reflect cellular generations. As far as we know, the rate of tissue turn-over in planarians has not been measured quantitatively. We defined *g* as the unknown ratio of cellular generation / organismal generation and evaluated its effect on the calculated *N*_*e,sc*_. In addition to the combined gen II and gene XII samples, we calculated *F*_*k*_ and *N*_*e*_ from all pairs of gen. II and gen. XII replicates to evaluate the effects of experimental variance (vs. biological variance or drift) on our findings.

Nucleotide diversity (*π*) for each sample was calculated from positions covered 10-60X in that sample as:

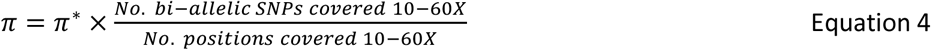

Where *π*^∗^ is nucleotide diversity at bi-allelic SNPs covered 10-60X and is calculated from the number of reference and alternative allele counts (*Rc* and *Ac*, respectively) as follows:

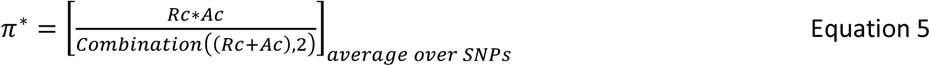

The second term on the right side of equation 4 is the proportion of polymorphic positions. No restriction on initial AAF was imposed for calculation of *π*.

The somatic mutation rate under the assumption of mutation-drift equilibrium was estimated according to the neutral theory expectations in haploid form:

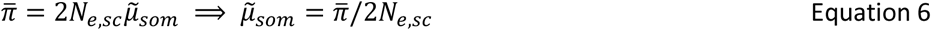

The haploid form of the neutral equation was chosen because in our Pool-seq data each genomic position in a sample is represented by individually sequenced chromosomes rather than diploid genotypes. Correspondingly, sample size at each position is set equal to the sequencing coverage at that position.

We also estimated *N*_*e*_ and somatic mutation rate by a second approach. We derived a new formula to model trans-generational change in heterozygosity (*π*) under mutation-drift non-equilibrium:

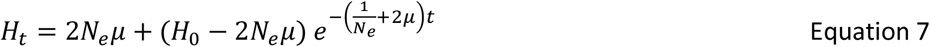

Details of the derivation are presented below. We used a nonlinear least-square fitting algorithm (algorithm “port” implemented in the R function nls()) to estimate *H*_0_(*π*_0_), *N*_*e*_ and *μ* from the rate of reduction in *π* over generations. *N*_*e*_ = 500, *μ* = 1 × 10^−5^, *π*_0_ = 0.005 where given as starting values and lower bound for all three was set to 0. Maximum iteration was set to 100.

### Deriving the equation for change of heterozygosity under mutation-drift non-equilibrium

The probability of identity by descent (IBD) designated *F* (sometimes called the inbreeding coefficient) equals relative loss of heterozygosity (*H*).

Note: This *F* has nothing to do with the *F*_*k*_ parameter introduced before in equations 1-3. It is a completely different entity.

In a diploid population, *F* is increased by drift and reduced by mutation according to the following equation:

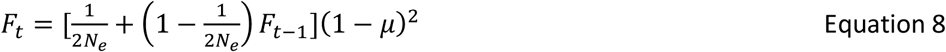

Under mutation-drift equilibrium *F*_*t*_ = *F*_*t*−1_; assuming *μ* ≪ 1, this reduces to:

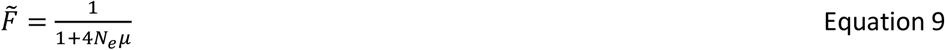

And:

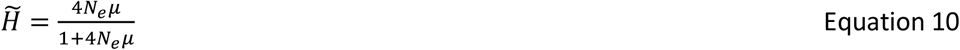

This result for ^*H*:^ is equal to the expected value of nucleotide diversity (*π*) under the infinite sites model. For small values of *π*, ^*H*:^ = *E*(*π*) ≅ 4*N*_*e*_*μ*.

These are classic results found in most population genetics text books^43,44^. But how can the recursive equation 11 be solved if equilibrium cannot be assumed? We found two solutions with equivalent results:

Solution 1: A trick that usually works in solving recursive equations finally reaching an equilibrium value is subtracting the equilibrium value from the two sides (of Eq. 11):

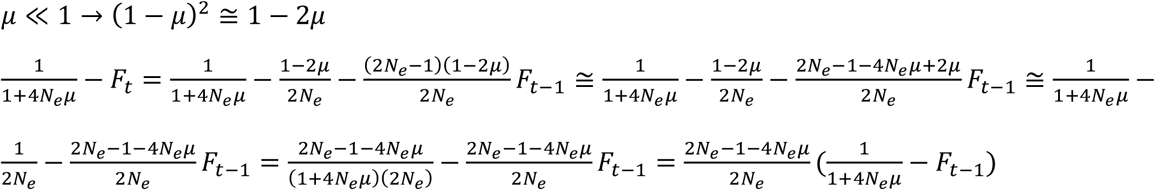

Now, we have a recursive that can be easily solved and taken back to *F*_0_:

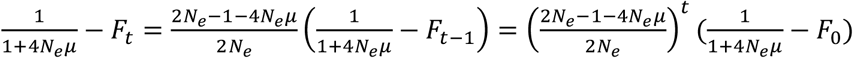

This simplifies to:

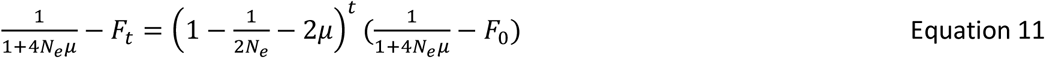

This means that the deviation of existing *F* from the equilibrium *F*: is reduced by a factor of 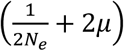 every generation. Solving for *H* and assuming 4*N*_*e*_*μ* ≪ 1:

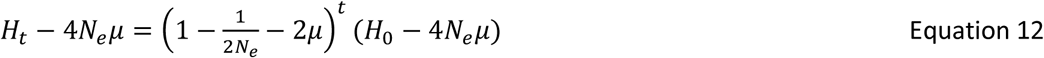

Solution 2: We can write the difference in *H* from generation to the next, convert discrete time to continuous time to arrive at a differential equation, and solve it. The difference equation can be easily obtained from eq. 11 but is also given in population genetics texts^44^:

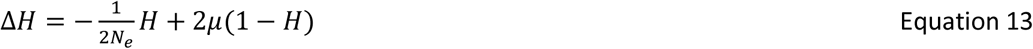

This is very intuitive: Heterozygosity is reduced by 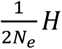 per generation as a result of drift, and is increased by 2*μ*(1 − *H*) via new mutations. The differential form of eq. 16 is:

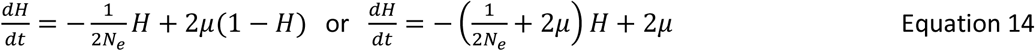

To general form of equation 17 and its integral are:

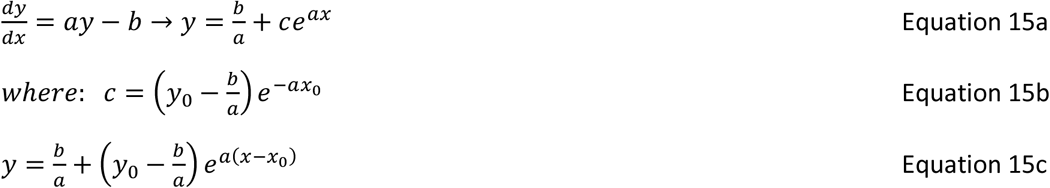

Substituting *y* with *H* and *x* with *t*:

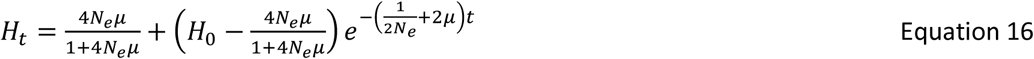

Assuming *π*: ≪ 1 like before:

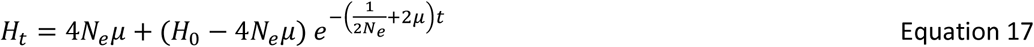

Noting 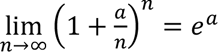, it is evident that for small 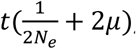, eq. 17 approximates eq. 12. We substituted 2*N*_*e*_ with *N*_*e*_ to be compatible with the haploid model we used throughout this study when applying this equation to the planaria data (Equation 7).

## Acknowledgement

This work was supported by NIH Grant No. GM098741, American Cancer Society Grant No. 130920-RSG-17-114-01-RMC and the postdoctoral PBBR grant by Sandler Foundation and UCSF.

---

**Table S1.** Accession number of Tricladida COI barcode sequences used for taxonomical identification of specimens and the number of reads from two representative samples from generations VI and X mapping to each barcode.

| Accession | Species | Sample 1 (gen. VI) | Sample 2 (gen. X) |
| --- | --- | --- | --- |
| GBPL4280-15 | *Girardia sp. AW2014* | 411 | 561 |
| GBPL1167-09 | *Girardia tigrina* | 298 | 504 |
| GBPL2853-13 | *Obama carinata* | 12 | 57 |
| GBPL2870-13 | *Choeradoplana bocaina* | 5 | 23 |
| GBPL2825-13 | *Pasipha tapetilla* | 3 | 13 |
| GBPL2856-13 | *Notogynaphallia plumbea* | 1 | 9 |
| GBPL2926-13 | *Luteostriata graffi* | 0 | 5 |
| GBPL2839-13 | *Luteostriata ceciliae* | 1 | 3 |
| GBPL1552-13 | *Obama ladislavii* | 1 | 2 |
| GBMAB2254-15 | *Platydemus manokwari* | 1 | 2 |
| GBPL2921-13 | *Endeavouria septemlineata* | 0 | 2 |
| GBPL2834-13 | *Choeradoplana gladysmariae* | 0 | 1 |
| GBPL2886-13 | *Issoca jandaia* | 0 | 1 |
| GBPL2907-13 | *Issoca rezendei* | 0 | 1 |
| GBPL3444-14 | *Luteostriata abundans* | 0 | 1 |
| GBPL2861-13 | *Notogynaphallia parca* | 0 | 1 |
| GBPL3196-14 | *Cephaloflexa bergi* | 1 | 0 |
| GBPL1559-13 | *Geoplana quagga* | 1 | 0 |
| GBPL2859-13 | *Pasipha chimbeva* | 1 | 0 |
| GBPL1173-09 | *Artioposthia testacea* | 0 | 0 |
| GBPL1172-09 | *Bipalium adventitium* | 0 | 0 |
| GBPL2841-13 | *Cephaloflexa araucariana* | 0 | 0 |
| GBPL2833-13 | *Choeradoplana albonigra* | 0 | 0 |
| GBPL2858-13 | *Choeradoplana banga* | 0 | 0 |
| GBPL1551-13 | *Choeradoplana iheringi* | 0 | 0 |
| GBPL2837-13 | *Cratera crioula* | 0 | 0 |
| GBPL2903-13 | *Cratera pseudovaginuloides* | 0 | 0 |
| GBPL2901-13 | *Cratera tamoia* | 0 | 0 |
| GBPL1169-09 | *Dendrocoelum lacteum* | 0 | 0 |
| GBPL1458-13 | *Dugesia aethiopica* | 0 | 0 |
| GBPL2668-13 | *Dugesia ariadnae* | 0 | 0 |
| GBPL1478-13 | *Dugesia benazzii* | 0 | 0 |
| GBPL2669-13 | *Dugesia cretica* | 0 | 0 |
| GBPL1472-13 | *Dugesia etrusca* | 0 | 0 |
| GBPL1469-13 | *Dugesia gonocephala* | 0 | 0 |
| GBPL1468-13 | *Dugesia hepta* | 0 | 0 |
| GBPL1467-13 | *Dugesia ilvana* | 0 | 0 |
| GBMTG2496-16 | *Dugesia japonica* | 0 | 0 |
| GBPL1464-13 | *Dugesia liguriensis* | 0 | 0 |
| GBPL1463-13 | *Dugesia notogaea* | 0 | 0 |
| GBPL1443-13 | *Dugesia sicula* | 0 | 0 |
| GBPL1460-13 | *Dugesia subtentaculata* | 0 | 0 |
| GBPL3422-14 | *Enterosyringa pseudorhynchodemus* | 0 | 0 |
| GBPL2927-13 | *Geobia subterranea* | 0 | 0 |
| GBPL2830-13 | *Geoplana chita* | 0 | 0 |
| GBPL2865-13 | *Geoplana goetschi* | 0 | 0 |
| GBPL2894-13 | *Geoplana hina* | 0 | 0 |
| GBPL2850-13 | *Geoplana pulchella* | 0 | 0 |
| GBPL2893-13 | *Geoplana vaginuloides* | 0 | 0 |
| GBPL1071-09 | *Girardia anderlani* | 0 | 0 |
| GBPL2828-13 | *Gusana* | 0 | 0 |
| GBPL1546-13 | *Imbira guaiana* | 0 | 0 |
| GBPL1549-13 | *Imbira marcusii* | 0 | 0 |
| GBPL2847-13 | *Luteostriata ernesti* | 0 | 0 |
| GBPL2849-13 | *Luteostriata muelleri* | 0 | 0 |
| GBPL2902-13 | *Matuxia tuxaua* | 0 | 0 |
| GBSP4320-12 | *Microplana robusta* | 0 | 0 |
| GBSP4311-12 | *Microplana terrestris* | 0 | 0 |
| GBPL2898-13 | *Notogynaphallia sexstriata* | 0 | 0 |
| GBPL1547-13 | *Obama burmeisteri* | 0 | 0 |
| GBPL2842-13 | *Obama josefi* | 0 | 0 |
| GBPL2826-13 | *Paraba franciscana* | 0 | 0 |
| GBPL2860-13 | *Paraba multicolor* | 0 | 0 |
| GBPL2869-13 | *Paraba phocaica* | 0 | 0 |
| GBMAB314-15 | *Parakontikia ventrolineata* | 0 | 0 |
| GBPL2889-13 | *Pasipha pasipha* | 0 | 0 |
| GBPL2891-13 | *Pasipha pinima* | 0 | 0 |
| GBPL2913-13 | *Pasipha rosea* | 0 | 0 |
| TERIN001-17 | *Rhynchodemus sylvaticus* | 0 | 0 |
| GBMTG5013-16 | *Schmidtea mediterranea* | 0 | 0 |
| GBPL1435-13 | *Schmidtea polychroa* | 0 | 0 |
| GBPL2827-13 | *Supramontana irritata* | 0 | 0 |

**Table S2.**
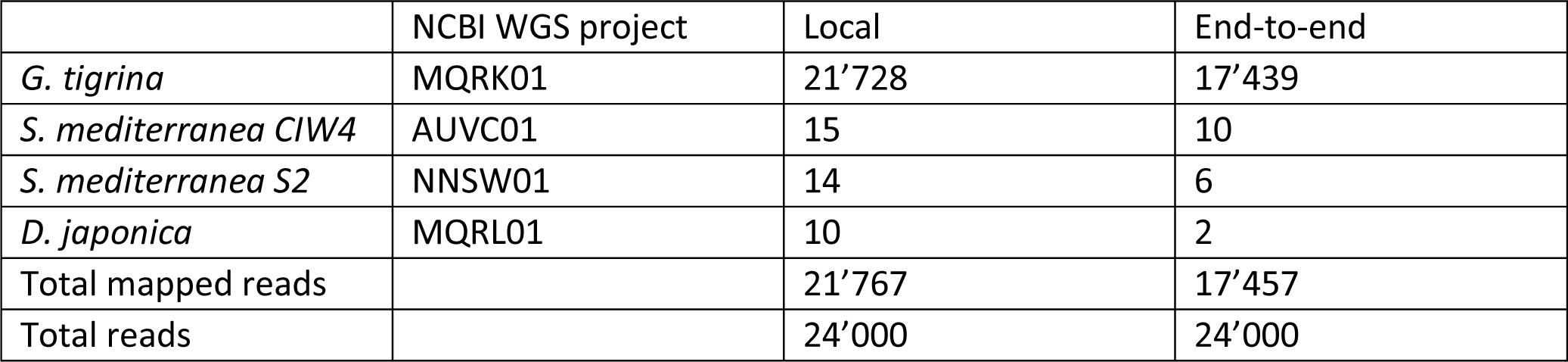
Subsampled-and-pooled reads mapping to the four Dugesiidae genomes.

|  | NCBI WGS project | Local | End-to-end |
| --- | --- | --- | --- |
| *G. tigrina* | MQRK01 | 21'728 | 17'439 |
| *S. mediterranea CIW4* | AUVC01 | 15 | 10 |
| *S. mediterranea S2* | NNSW01 | 14 | 6 |
| *D. japonica* | MQRL01 | 10 | 2 |
| Total mapped reads |  | 21'767 | 17'457 |
| Total reads |  | 24'000 | 24'000 |

**Table S3.** Details of quality control and mapping statistics of trimmed and deduplicated fastq files. The R1 fastq file from sequencing lane 8 of the generation 14 (sample XIV_L008) was damaged during a server transfer and could not be recovered. This sample was pre-processed like others but as a single-end sequenced library. The Raw fastqc files had varying levels of low quality read segments (especially towards the 3’ end and more in the reverse read or 2^nd^ mate files) and PCR duplicates. They did not contain adapter or any other over-represented sequences (other than N homopolymers in some cases). Across the 12 paired-end samples, the fractions of proper pairs after processing by Trimmomatic and Clumpify were 87-95% and 67-85%, respectively. Clumpify was very high-memory-consuming and worked for all samples only if the -Xmx option were set to 230g or higher. Min. deduplication (%): “percent of sequences remaining if deduplicated” from the output of FastQC after running Clumpify. This is a minimum estimate because it is based on read sequence identity inferred separately from forward and reverse files while Clumpify identifies duplication based on both mates of a pair. Mapping rate: percentage of trimmed and deduplicated read pairs that were mapped to the *G. tigrina* reference genome by *bwa mem*.

| Sample | Raw pairs | Trimmomatic output pairs | Clumpify output pairs | Min. deduplication (%) | Mapping rate (proper pairs) (%) |
| --- | --- | --- | --- | --- | --- |
| II_L003 | 179'975'602 | 171'531'579 | 152'492'295 | 84.93; 77.74 | 96.22 (73.99) |
| II_L005 | 172'108'843 | 163'149'585 | 144'494'228 | 85.36; 77.76 | 96.23 (74.11) |
| II_L006 | 178'680'161 | 169'213'772 | 150'056'408 | 85.93; 78.82 | 96.20 (74.16) |
| VI_L002 | 200'302'395 | 189'399'008 | 149'305'526 | 84.83; 78.15 | 93.90 (70.94) |
| VI_L003 | 196'579'360 | 171'891'056 | 141'505'349 | 81.85; 80.74 | 92.81 (68.76) |
| VIII_L005 | 191'266'046 | 166'653'906 | 130'689'674 | 83.56; 83.9 | 93.19 (68.04) |
| VIII_L006 | 191'085'892 | 166'682'020 | 130'303'177 | 82.45; 80.27 | 93.47 (68.48) |
| X_L007 | 186'274'991 | 167'125'904 | 127'102'268 | 82.44; 80.46 | 92.81 (69.80) |
| X_L008 | 188'302'804 | 168'925'128 | 128'794'306 | 82.71; 81.52 | 92.82 (69.77) |
| XII_L005 | 184'648'701 | 165'342'241 | 130'904'629 | 81.48; 76.55 | 92.66 (69.01) |
| XII_L006 | 192'481'056 | 170'964'293 | 129'036'162 | 79.8; 80.61 | 92.17 (68.30) |
| XIV_L007 | 197'049'701 | 183'302'121 | 152'225'064 | 82.87; 82.2 | 92.64 (68.56) |
| XIV_L008* | 190'083'700 | 169'937'444 | 110'184'051 | 92.09 | 84.80 (N/A) |

\* Only the reserve file was available from this sample. All numbers presented for this row represent single reads not pairs.

**Table S4.**
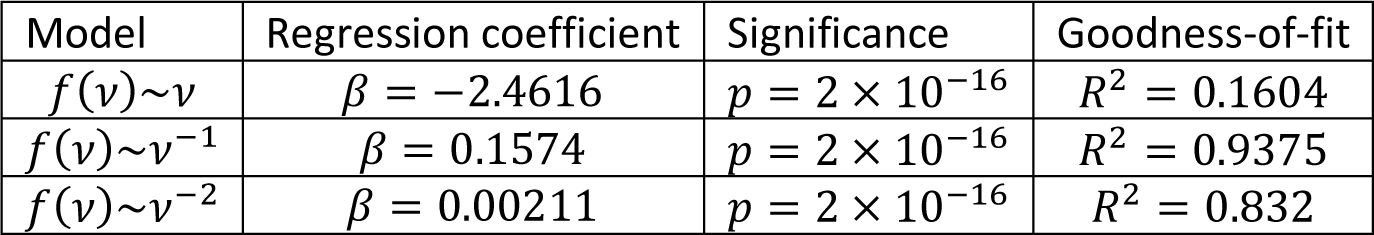
Evaluating the goodness-of-fit of AFS with neutrality (drift) vs. strong linked selection (draft). Positions with derived allele frequency *ν*: 0.01 − 0.9 were included to avoid extreme outlier effects and fixed substitutions. Variables *ν*. *inv* = *ν*^−1^ and *ν*. *inv*. *sq* = *ν*^−2^ were created and the linear fit of *f*(*ν*) against them was examined. The simple *f*(*ν*)∼*ν* linear model was tested as a known bad fit (negative control).

| Model | Regression coefficient | Significance | Goodness-of-fit |
| --- | --- | --- | --- |
| $f(\nu) \sim \nu$ | $\beta = -2.4616$ | $p = 2 \times 10^{-16}$ | $R^2 = 0.1604$ |
| $f(\nu) \sim \nu^{-1}$ | $\beta = 0.1574$ | $p = 2 \times 10^{-16}$ | $R^2 = 0.9375$ |
| $f(\nu) \sim \nu^{-2}$ | $\beta = 0.00211$ | $p = 2 \times 10^{-16}$ | $R^2 = 0.832$ |

**Table S5.** Evaluating the reproducibility of *F̅*_*k*_ and *N*_*e*_ estimates from unmerged replicates of generations II and XII (assuming *t* = 10). The corresponding values from the merged-rep calculation were *F̅*_*k*_ = 0.1433 and *N*_*e*_ = 141.70. This tables demonstrates the magnitude of variation that results from experimental stochasticity among replicates. There is a substantial (near three-fold) change in the estimates of*N*; but the order of magnitude remains in the range of 10^2^s.

| $Gen_i$ | $Gen_j$ | $S_0$ | $S_t$ | $\bar{F}_k$ | $N_e$ |
| --- | --- | --- | --- | --- | --- |
| II.3 | XII.5 | 20.21916 | 16.54553 | 0.177164 | 135.1165 |
| II.5 | XII.5 | 19.53791 | 16.54553 | 0.172657 | 149.3833 |
| II.6 | XII.5 | 21.18524 | 16.54553 | 0.172808 | 139.6204 |
| II.3 | XII.6 | 20.21916 | 16.80986 | 0.140765 | 291.0595 |
| II.5 | XII.6 | 19.53791 | 16.80986 | 0.125863 | 614.5872 |
| II.6 | XII.6 | 21.18524 | 16.80986 | 0.126612 | 467.6062 |

**Figure S1.**
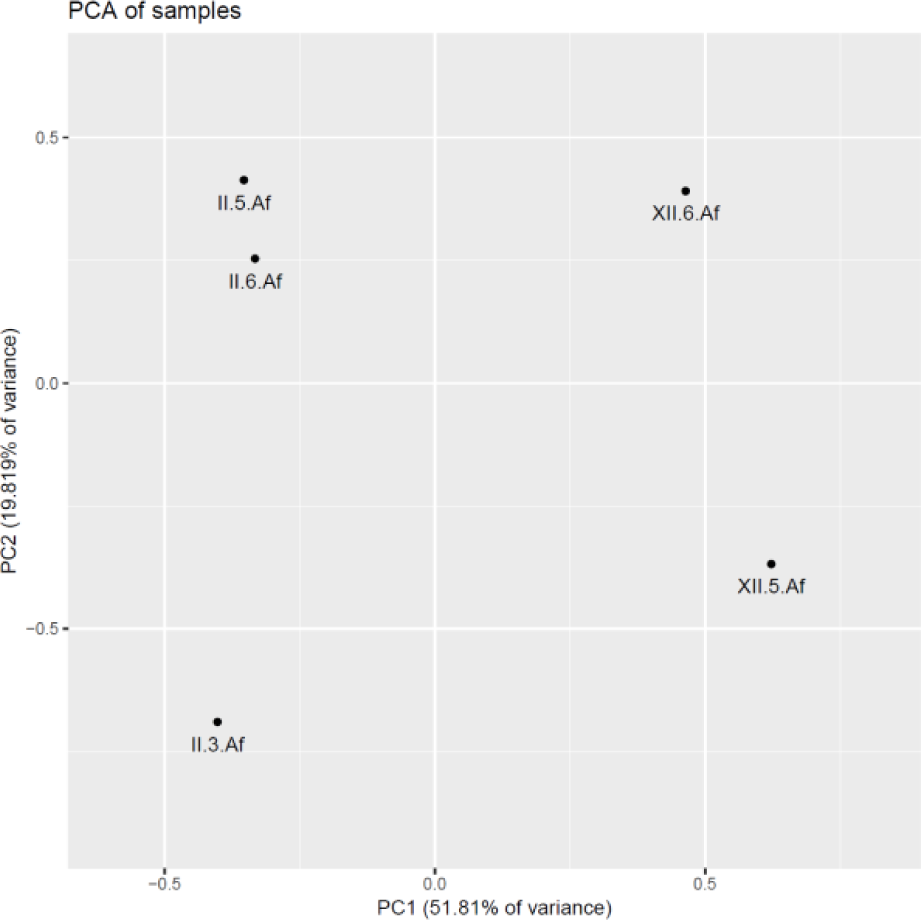
PCA of allele frequencies of replicates from samples II and XII. PC1 which explains ∼52% of the variance (and more than twice as much as PC2) clearly separates II.3, II.5 and II.6 from XII.5 and XII.6. This confirms that variation due to drift is essentially larger than technical variability.

**Figure S2.**
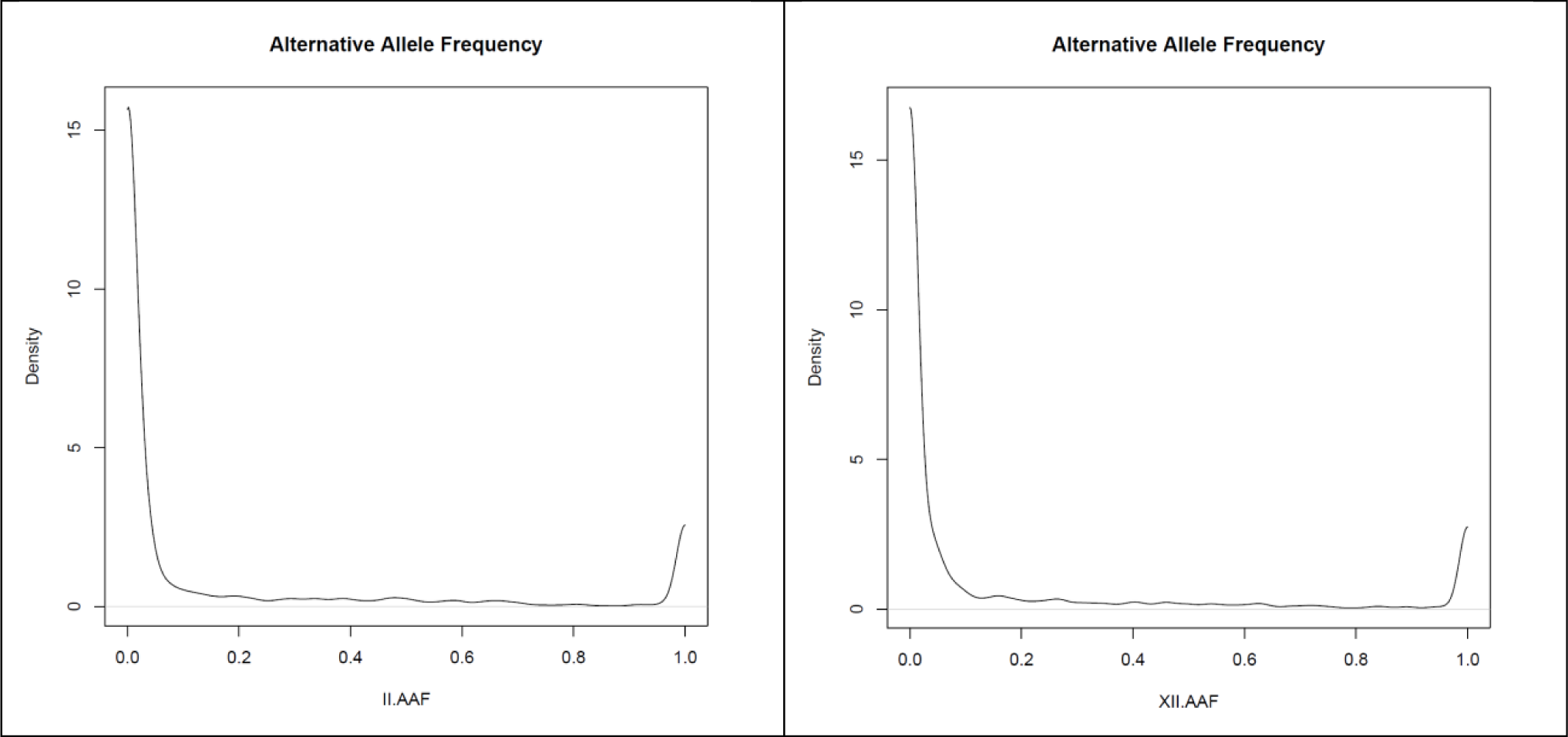
Allele frequency spectra of merged-rep samples II and XII. Positions with *f*_*alt*_ = 1 would not be normally included in the AFS because they are not polymorphic. They are recorded here because we ran FreeBayes with the --use-reference-allele option. They provide a visual comparison of divergence (fixed substitutions) from the reference versus segregating variation. AAF: alternative allele frequency.

**Figure S3.**
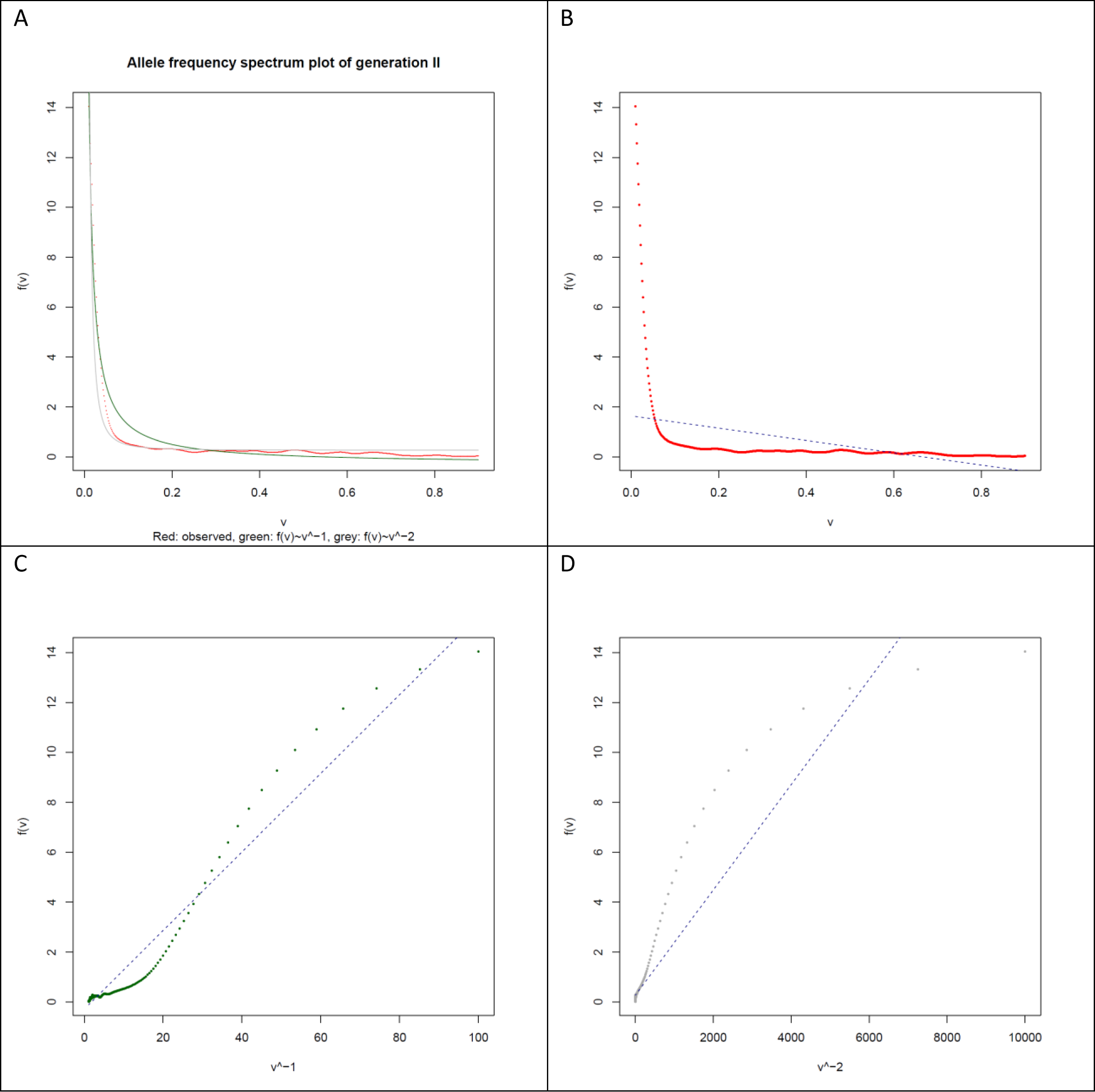
Goodness of fit of the allele frequency spectrum with drift and draft models. AFS (shown for merged-rep sample II here) is most compatible with the *f*(*v*)∼*v*^−1^ function which suggests that drift (not selection) is the strongest force shaping it. A: Observed allele frequency density *f*(*v*) plotted against allele frequency *v* along with predictions of drift (green) and draft (grey) models. Note that the measure of goodness-of-fit is the vertical distance between observed and expected. B, C and D: Visualizing the goodness-of-fit of the best linear models describing *f*(*v*) as a function of *v* (B), *v*^−1^ (C) and *v*^−2^ (D). Although none of the three shows a perfect fit, *f*(*v*)∼*v*^−1^ is a better fit than the other two.

**Figure S4.**
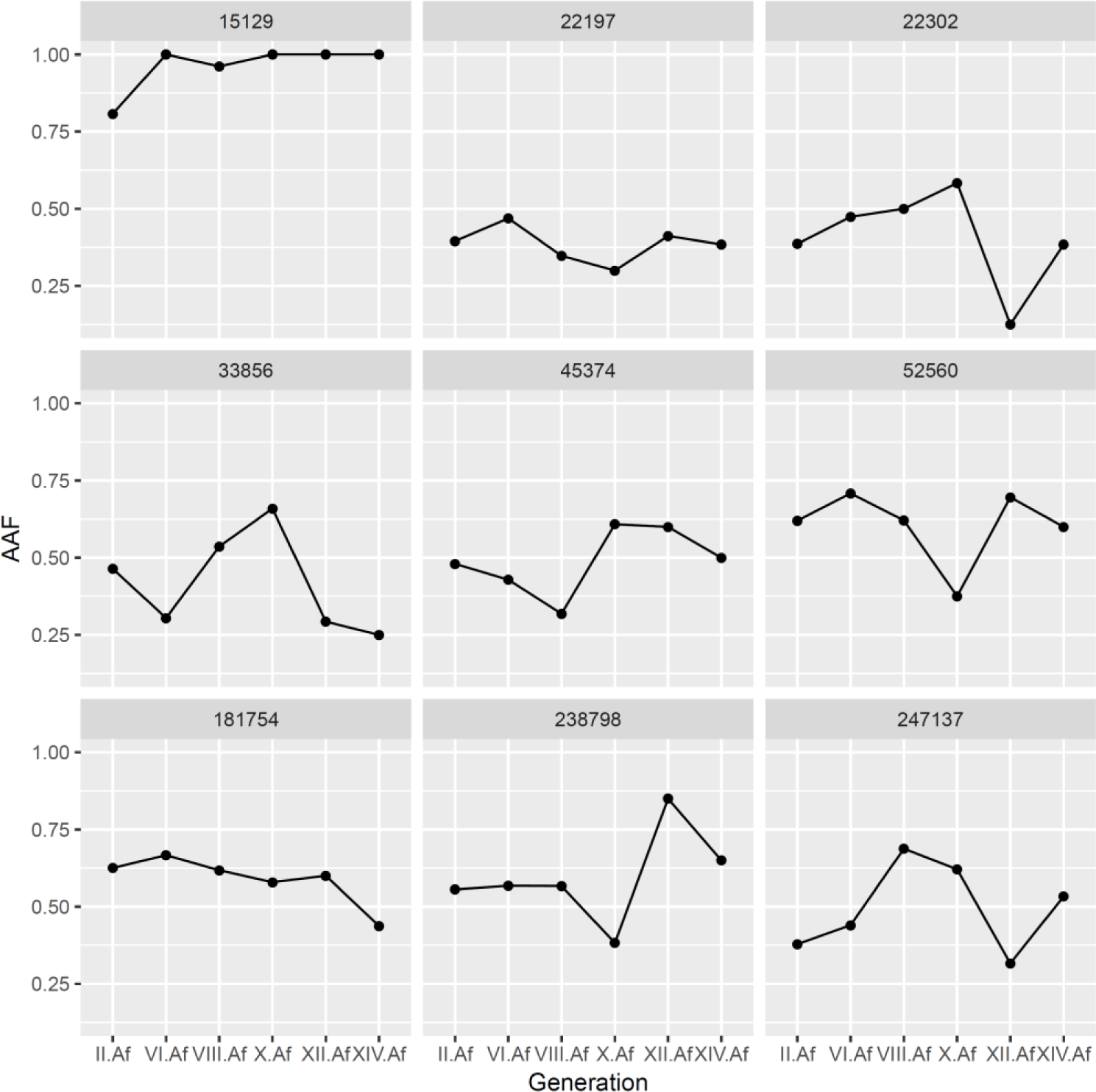
Transgenerational fluctuation of AAF (alternative allele frequency) in 9 randomly selected SNPs with AAF 0.1-0.9 at generation II. The same criterion was applied to include SNPs in calculation of *N*_*e,sc*_. All positions were covered 10-60X in all samples. The observed changes in AAF are due to somatic drift between generations + sampling variance at each generation. Number on top of panels: position on scaffold MQRK01218062.1.

**Figure S5.**
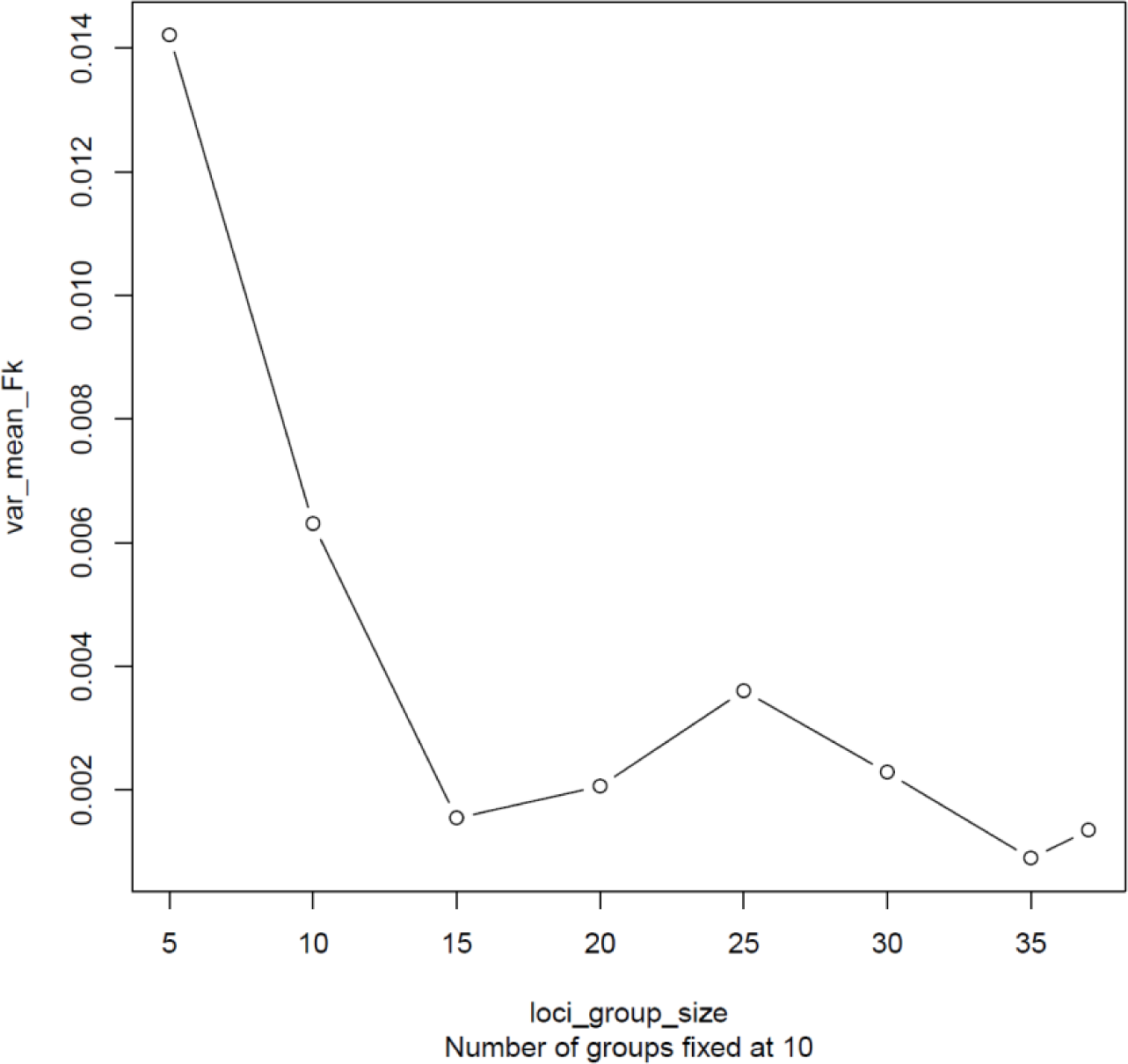
Evaluating the independence of SNPs through the variance of their *F*_*k*_ estimates. The SNPs were subsampled (without replacement) into 10 groups of variable sizes (*n* = 5, 10, 15, 20, 25, 30, 35). Within group means (*F̄*_*k*_s) were calculated and then *var*(*F̄*_*k*_) across groups was obtained. *var*(^*F̄*^_*k*_) decreases going from *n* = 5 to *n* = 15 and plateaus after. This seems consistent with the loci providing at least partially independent information.

## References

1. Rink, J. C. Stem cell systems and regeneration in planaria. Dev. Genes Evol. 223, 67–84 (2013).

2. Reddien, P. W. & Alvarado, A. S. Fundamentals of Planarian Regeneration. Annu. Rev. Cell Dev. Biol. 20, 725–757 (2004).

3. Knakievicz, T., Vieira, S. M., Erdtmann, B. & Ferreira, H. B. Reproductive modes and life cycles of freshwater planarians (Platyhelminthes, Tricladida, Paludicula) from southern Brazil. Invertebr. Biol. 125, 212–221 (2006).

4. Lázaro, E. M. et al. Molecular barcoding and phylogeography of sexual and asexual freshwater planarians of the genus *Dugesia* in the Western Mediterranean (Platyhelminthes, Tricladida, Dugesiidae). Mol. Phylogenet. Evol. 52, 835–845 (2009).

5. Reddien, P. W. The Cellular and Molecular Basis for Planarian Regeneration. Cell 175, 327–345 (2018).

6. Lopes, K. A. R., Velho, N. M. R. D. C. & Soares, C. P. Method of isolation and characterization of *Girardia tigrina* stem cells. Biomed. Reports 3, 163–166 (2015).

7. Isaeva, V., Alexandrova, Y. & Reunov, A. Interaction between chromatoid bodies and mitochondria in neoblasts and gonial cells of the asexual and spontaneously sexualized planarian, *Girardia (Dugesia) tigrina*. Invertebr. Reprod. Dev. 48, 119–128 (2005).

8. Newmark, P. A. & Alvarado, A. S. Not Your Father’S Planarian: a Classic Model Enters the Era of Functional Genomics. Nat. Rev. Genet. 3, 210–219 (2002).

9. Zeng, A. et al. Prospectively Isolated Tetraspanin + Neoblasts Are Adult Pluripotent Stem Cells Underlying Planaria Regeneration. Cell 173, 1593–1608.e20 (2018).

10. Nishimura, O. et al. Unusually large number of mutations in asexually reproducing clonal planarian *Dugesia japonica*. PLoS One 10, 1–23 (2015).

11. Otto, S. P. & Orive, M. E. Evolutionary consequences of mutation and selection within an individual. Genetics 141, 1173–1187 (1995).

12. Otto, S. P. & Hastings, I. M. Mutation and selection within the individual. Genetica 102–103, 507– 524 (1998).

13. Gehrke, A. R. & Srivastava, M. Neoblasts and the evolution of whole-body regeneration. Curr. Opin. Genet. Dev. 40, 131–137 (2016).

14. Rozanski, A. et al. PlanMine 3.0-improvements to a mineable resource of flatworm biology and biodiversity. Nucleic Acids Res. 1–9 (2018). doi:10.1093/nar/gky1070

15. Scimone, M. L., Kravarik, K. M., Lapan, S. W. & Reddien, P. W. Neoblast specialization in regeneration of the planarian *Schmidtea mediterranea*. Stem Cell Reports 3, 339–352 (2014).

16. Neher, R. A. & Shraiman, B. I. Genetic draft and quasi-neutrality in large facultatively sexual populations. Genetics 188, 975–996 (2011).

17. Knezović, L., Miliša, M., Kalafatić, M., Rajević, N. & Planinić, A. A key to the freshwater triclads (Platyhelminthes, Tricladida) of Herzegovina watercourses. Period. Biol. 117, 425–433 (2015).

18. Waples, R. S. A generalized approach for estimating effective population size from temporal changes in allele frequency. Genetics 121, 379–91 (1989).

19. Jorde, P. E. & Ryman, N. Temporal allele frequency change and estimation of effective size in populations with overlapping generations. Genetics 139, 1077–1090 (1995).

20. Berthier, P. et al. Likelihood-based estimation of the effective population size using temporal changes in allele frequencies: a genealogical approach. Genetics 160, 741–51 (2002).

21. Waples, R. S. & Yokota, M. Temporal estimates of effective population size in species with overlapping generations. Genetics 175, 219–233 (2007).

22. Lobo, D., Beane, W. S. & Levin, M. Modeling planarian regeneration: A primer for reverse-engineering the worm. PLoS Comput. Biol. 8, (2012).

23. Baguñà, J., Saló, E. & Auladell, C. Regeneration and pattern formation in planarians \nIII. Evidence that neoblasts are totipotent stem cells and the source of blastema cells. Development 107, 77– 86 (1989).

24. Elliott, S. A. & Sánchez Alvarado, A. The history and enduring contributions of planarians to the study of animal regeneration. Wiley Interdiscip. Rev. Dev. Biol. 2, 301–326 (2013).

25. Montgomery, J. R. & Coward, S. T. On the Minimal Size of a Planarian Capable of Regeneration. Trans. Am. Microsc. Soc. 93, 386–391 (1974).

26. Pollak, E. A New Method for Estimating the Effective Population Size from Allele Frequency Changes. Genetics 104, 531–548 (1983).

27. Guo, L., Zhang, S., Rubinstein, B., Ross, E. & Alvarado, A. S. Widespread maintenance of genome heterozygosity in *Schmidtea mediterranea*. *Nat*. Ecol. Evol. 1, 0019 (2016).

28. Pellettieri, J. Regenerative tissue remodeling in planarians – The mysteries of morphallaxis. Semin. Cell Dev. Biol. 87, 13–21 (2019).

29. Nicolaisen, L. E. & Desai, M. M. Distortions in Genealogies due to Purifying Selection and Recombination. Genetics 195, 221–230 (2013).

30. Charlesworth, B. Effective population size and patterns of molecular evolution and variation. Nat. Rev. Genet. 10, 195–205 (2009).

31. Campbell, P. J. & Martinocorena, I. Somatic mutation in cancer and normal cells. Science (80-.). 349, 1483–1489 (2015).

32. Lynch, M. Evolution of the mutation rate. Trends Genet. 26, 345–352 (2010).

33. Sniegowski, P. D., Gerrish, P. J., Johnson, T. & Shaver, A. The evolution of mutation rate: separating causes from consequences. BioEssays 22, 1057–1066 (2000).

34. Andre, J. B. & Godelle, B. The evolution of mutation rate in finite asexual populations. Genetics 172, 611–626 (2006).

35. Dunham, J. P. & Friesen, M. L. A cost-effective method for high-throughput construction of Illumina sequencing libraries. Cold Spring Harb. Protoc. 9, 820–834 (2013).

36. Bolger, A. M., Lohse, M. & Usadel, B. Trimmomatic: A flexible trimmer for Illumina sequence data. Bioinformatics 30, 2114–2120 (2014).

37. Hebert, P. D. N., Cywinska, A., Ball, S. L. & deWaard, J. R. Biological identifications through DNA barcodes. Proc. Biol. Sci. 270, 313–21 (2003).

38. Kofler, R., Langmuller, A. M., Nouhaud, P., Otte, K. A. & Schlotterer, C. Suitability of Different Mapping Algorithms for Genome-wide Polymorphism Scans with Pool-Seq Data. G3 6, 3507–3515 (2016).

39. Garrison, E. & Marth, G. Haplotype-based variant detection from short-read sequencing. 1–9 (2012).

40. Hahn, C., Fromm, B. & Bachmann, L. Comparative genomics of flatworms (Platyhelminthes) reveals shared genomic features of ecto- and endoparastic neodermata. Genome Biol. Evol. 6, 1105–1117 (2014).

41. Grohme, M. A. et al. The genome of *Schmidtea mediterranea* and the evolution of core cellular mechanisms. Nature 554, 56–61 (2018).

42. Waples, R. S. Temporal Variation in Allele Frequencies: Testing the Right Hypothesis. Evolution (N. Y). 43, 1236–1251 (1989).

43. Crow, J. F. & Kimura, M. An Introduction to population genetics theory. (Harper and Row, 1970).

44. Gillespie, J. H. Population genetics, a concise guide. (The John Hopkins University Press, 2004).

